# Acute exercise increases systemic kynurenine pathway metabolites and activates the AHR in human PBMCs

**DOI:** 10.1101/2024.01.17.576018

**Authors:** Niklas Joisten, David Walzik, Alexander Schenk, Alan J Metcalfe, Sergen Belen, Kirill Schaaf, Sebastian Gehlert, Polyxeni Spiliopoulou, Ann-Marie Garzinsky, Mario Thevis, Ludwig Rappelt, Lars Donath, Sven G Meuth, Wilhelm Bloch, Philipp Zimmer

## Abstract

The kynurenine pathway of tryptophan degradation generates several metabolites such as kynurenine or kynurenic acid that serve as endogenous ligands of the aryl hydrocarbon receptor (AHR). Due to its distinct biological roles particularly modulating the immune system, the AHR is a current therapeutic target across different inflammation-related diseases. Here, we show an exercise-induced increase in AHR ligand availability on a systemic level and a kynurenine pathway activation in peripheral mononuclear blood cells (PBMCs). Concurrently, the AHR is activated in PBMCs following acute exercise, with effects being dependent on exercise intensity. In conclusion, these data indicate a novel mechanistic link how exercise modulates the immune system through the kynurenine pathway-AHR axis, potentially underlying exercise-induced benefits in various chronic diseases.

## Main text

Dysregulations in the kynurenine pathway (KP) of tryptophan metabolism is a common pathophysiological feature of many chronic diseases, including metabolic, neuropsychiatric, and autoimmune disorders^1,2^. Many KP metabolites serve systemically as signaling molecules for different biological processes, i.e., immunomodulation, neuromodulation, and redox state. Under pathological conditions, KP metabolic flux is upregulated and imbalanced, often but not always towards the NAD^+^ precursor quinolinic acid. In consequence of this imbalanced metabolic breakdown, the activity of KP metabolite target receptors, such as the aryl hydrocarbon receptor (AHR) and N-methyl-D-aspartate (NMDA) receptors, subsequently disequilibrates^3^. The manipulation of key enzymes of the KP (e.g., indoleamine 2,3-dioxygenase-1 (IDO-1), kynurenine 3-monooxygenase (KMO)) has therefore become a promising therapeutic strategy across different chronic diseases, including but not limited to oncological and neurological disorders^4^.

Besides pharmacological manipulation of key KP enzymes, physical exercise, as powerful tool for the prevention and treatment of numerous chronic diseases, has been described as a potent KP modulator in skeletal muscle and systemically. Agudelo et al.^5^ showed that exercised skeletal muscle increases the metabolic flux towards kynurenic acid (KA), thereby mediating resilience to stress-induced depression in mice. In humans, exercise-induced increases in systemic levels of KA were successfully translated and have been reported in response to different exercise modalities, with aerobic exercise appearing to be the most potent stimulus^6– 8^. The role of exercise intensity as an essential factor for various biological adaptions, however, remains unclear in this context.

Despite several studies showing systemic increases in KP metabolites in response to exercise, possible signal transduction via target receptors have not yet been addressed. One of the most prominent KP metabolite receptors that has comprehensive roles in regulating immune homeostasis and which is expressed in various immune cells is the AHR^9^. Of note, the AHR is expressed across many different human tissues and mediates both inflammatory and anti-inflammatory regulatory mechanisms, which often depends on the biological context (e.g., type of ligands, time of activation).^10^ For instance, the AHR is involved in the differentiation and polarization of T cell subpopulations. It has been described for CD4^+^ T-cells and CD4^+^ TH-17-cells that activation of the AHR mediates the (trans-)differentiation into anti-inflammatory, regulatory T cells (T_regs_)^11^. Regulatory functions of the AHR have also been described for other immune cell types including natural killer (NK)- and B-cells^12,13^. The AHR has therefore become a promising therapeutic target in oncological diseases and beyond and is already tested in phase 1 clinical trials in persons with advanced tumors (e.g., NCT04069026; NCT04999202). Considering that previous exercise studies consistently reported an increased systemic AHR ligand availability as indicated by exercise-induced elevations of kynurenic acid levels together with observation of an upregulated intracellular KP metabolism in peripheral blood mononuclear cells (PBMCs), we here investigated whether the AHR is activated in PBMCs in response to acute exercise. We further examined a potential intensity-dependent effect on systemic KP metabolite levels as well as on KP enzymes and AHR activation in PBMCs. In an experimental human randomized cross-over trial, 24 healthy adults (see table 1 for participant characteristics) conducted an acute bout of both high-intensity interval training (HIIT) and work-matched moderate-intensity continuous training (MICT) on separate days. Blood samples were collected under standardized conditions immediately before (pre, baseline), immediately after (post), and 1h after (1h post) exercise (Figure 1A).

**Table 1.** Participant characteristics.

|  | Overall (n=24) |  | Women (n=12) |  | Men (n=12) |  |
| --- | --- | --- | --- | --- | --- | --- |
|  | *M* | *SD* | *M* | *SD* | *M* | *SD* |
| Age (years) | 30 | 4.33 | 28 | 4.09 | 32 | 3.80 |
| Height (cm) | 176.96 | 8.26 | 171.17 | 3.90 | 182.75 | 7.36 |
| Weight (kg) | 69.70 | 11.17 | 61.91 | 6.93 | 77.50 | 8.96 |
| BMI (kg×m<sup>-2</sup>) | 22.20 | 2.38 | 21.05 | 2.28 | 23.36 | 1.94 |
| RPE<sub>max</sub> | 19 | 1 | 19 | 1 | 19 | 1 |
| VO<sub>2peak</sub> (mL×kg<sup>-1</sup>×min<sup>-1</sup>) | 56.64 | 6.43 | 54.42 | 4.13 | 58.86 | 7.65 |
| HR<sub>max</sub> (bpm) | 181 | 11 | 183 | 11 | 180 | 11 |

M = Mean; SD = standard deviation; RPE = rate of perceived exhaustion; VO<sub>2peak</sub> = peak oxygen consumption; BMI = body mass index; HR<sub>max</sub> = maximum heart rate; bpm = beats per minute.

**Figure 1.**
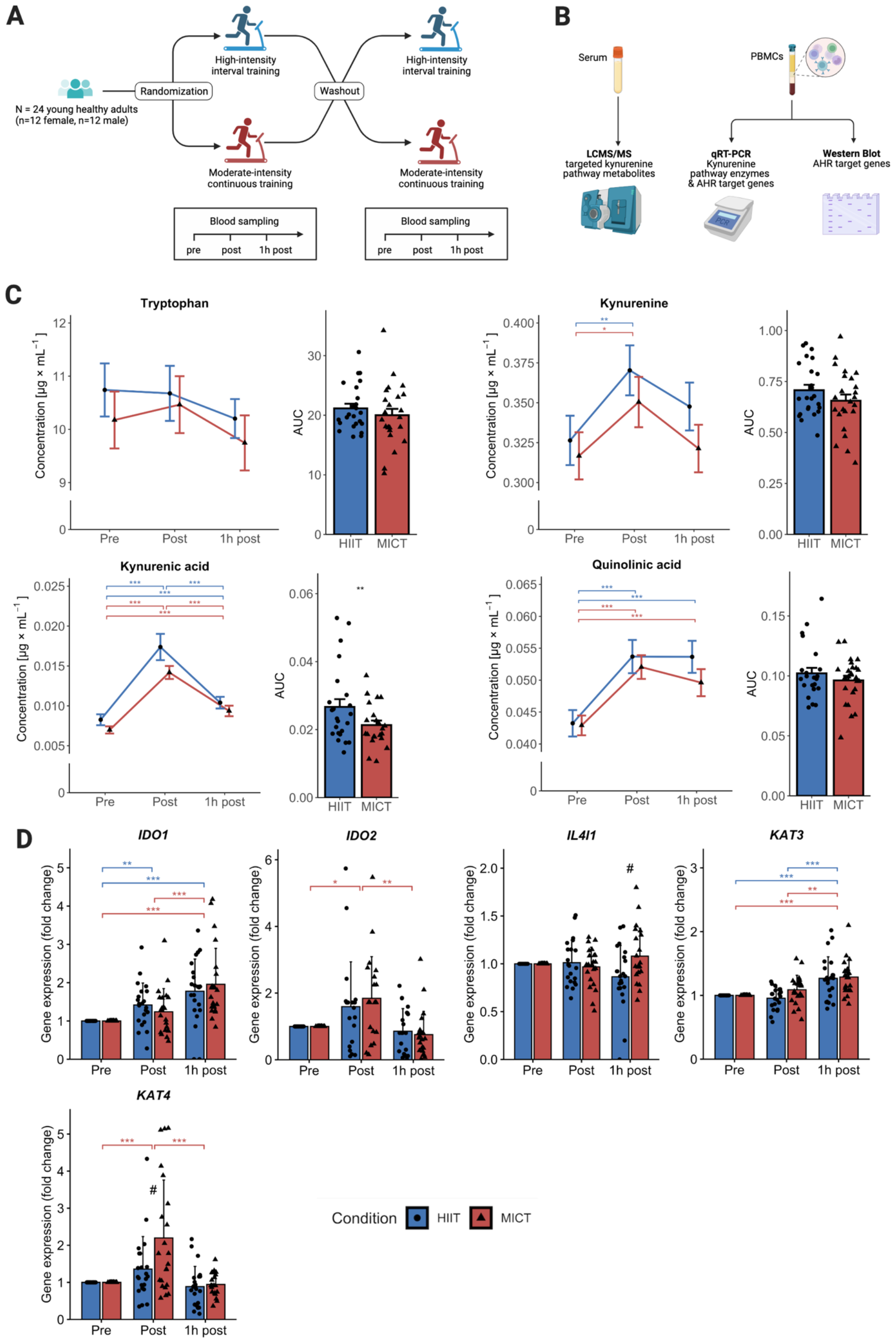
Study design and exercise-intensity dependent effects on levels of systemic kynurenine pathway metabolites. A: Schematic illustration of the randomized cross-over study design. B: Analysis plan of collected blood samples. C: Serum levels of tryptophan (HIIT: n=24; MICT: n=23) and its metabolites kynurenine (HIIT: n=24; MICT: n=23), kynurenic acid (HIIT: n=24; MICT: n=21), and quinolinic acid (HIIT: n=24; MICT: n=23) in response to acute high-intensity interval training (HIIT) and moderate-intensity continuous training (MICT). D: Gene expression levels of kynurenine pathway enzymes in PBMCs in response to acute HIIT (n=22) and MICT (n=22). Significances are based on analyses of variance (ANOVAs) with repeated measures and Bonferroni-corrected post hoc comparisons. Independent t-test was used for area under the curve (AUC) comparisons of HIIT vs. MICT. * p < .05; ** p < .001; *** p < .0001.

### Acute exercise increases systemic kynurenine pathway metabolites

In line with previous investigations^7,14^, acute exercise substantially altered systemic levels of tryptophan metabolites along the KP. While there was no significant change in tryptophan concentration, levels of kynurenine (KYN), KA, and quinolinic acid increased significantly in response to both exercise modalities (Figure 1C). The repeated measures ANOVA did not show a significant interaction effect between HIIT and MICT for any of the KP metabolites (supplement S3). However, when considering the area under the curve (AUC) as measure of ligand availability during the observed time frame (110 minutes in total), a greater AUC was observed for HIIT compared to MICT for KA (figure 1C). This finding suggests that, despite matching both exercise bouts for duration and energy demand, HIIT induces a greater transient increase in systemic KA levels (∼2-fold) compared to MICT, although the physiological relevance of this between-group difference remains questionable. Given that KA is one of the most potent endogenous AHR ligands^15,16^, an acute exercise-induced elevated AHR ligand availability in the circulation was detected in response to both HIIT and MICT. The systemic KA levels reached immediately post exercise are at least comparable or even higher than the concentration used in the benchmark paper that exposed KA as potent AHR ligand in cell culture experiments^16^.

### PBMCs show upregulated kynurenine pathway enzyme gene expression after exercise

It is well-established in rodents and humans that exercise mediates the expression of kynurenine-aminotransferases (KATs) in skeletal muscle, which is regulated by the transcriptional coactivator and master switch for energy metabolism PGC-1α1^5,6^. Whether the increases in systemic KA levels in response to acute exercise are solely generated by skeletal muscle or whether other source tissues might amplify KP metabolism remains unclear. Since many circulating immune cells are highly responsive to acute exercise and adjust distinct metabolic features upon activation^17,18^, we evaluated a potential exercise intensity-dependent effect on gene expression of key enzymes of the KP-AHR axis in PBMCs using qRT-PCR. We found that gene expression of all measured KP enzymes (*IDO1, IDO2, KAT3, KAT4*) was increased either immediately post exercise or 1h post exercise except for *IL4I1* (Figure 1D), suggesting intracellular metabolic KP activation by acute exercise in PBMCs. Gene expression of the recently described metabolic immune checkpoint *IL4I1*, which activates the AHR through generation of endogenous ligands including indoles and KA^15^, showed no effect over time. This finding suggests an inferior role of systemic indole levels in exercise-induced AHR activation. Considering the different exercise modes, MICT provoked greater gene expression compared to HIIT in *KAT4* (Figure 1D). Taken together, these data show that an activated KP metabolism by exercise is not only present in skeletal muscle but also in circulating immune cells. The contribution to exercise-induced systemic increases in KA by different source tissues is a matter of future investigations. However, an upregulated intracellular KP metabolism in response to acute exercise solidifies a potential AHR activation as target receptor for KYN and KA.

### Acute exercise activates the AHR in PBMCs in an intensity-dependent manner

To examine the impact of acute exercise on the AHR in PBMCs, we next assessed AHR gene expression as well as a comprehensive set of AHR target genes following HIIT and MICT. Both exercise modes significantly increased AHR expression (Figure 2A), indicating an AHR induction by acute exercise in PBMCs. Further, all AHR target genes, representing the surrogate parameter for AHR activation, increased significantly following both HIIT and MICT (Figure 2A). Particularly *CYP1B1* as one of the most described AHR targets substantially increased at 1h post exercise with remarkably consistent individual kinetics (Supplement file 2). Moreover, the AHR target gene *AHRR*, known for its negative feedback loop on AHR activation, was profoundly increased immediately post exercise, despite showing high inter-individual variability (Supplement file 2). To evaluate whether increased AHR target gene expression also affect their protein levels, we conducted western blot analysis of CYP1B1 and AHRR. While particularly CYP1B1 protein levels tended to increase, effects were not statistically significant and display broad inter-individual variability (Figure 2B). Discrepancies between mRNA and protein levels occur frequently^19^. In our study, the observation period might be too short to detect a consistent increase in protein levels. Albeit both exercise modes, HIIT and MICT, induce profound increases in AHR target genes, the effects driven by HIIT are significantly greater for *AHR* and *CYP1B1* compared to MICT (Figure 2A and Supplement 3). This finding is in accordance with greater systemic KA availability following HIIT, as previously reported (Figure 1C). In summary, our results show that acute exercise activates the AHR in PBMCs in an exercise-intensity dependent manner. In line with other prominent exerkines, such as IL-6, increased systemic KA levels return rapidly back to baseline levels after exercise cessation. Although the observed exercise-induced increase in AHR ligands (i.e. KA, KYN) is transient in nature, subsequent AHR activation in PBMCs may lead to relevant longer-term adaptions. Considering the transient increase in AHR ligands and that most biological effects induced by acute exercise typically occur within a 48-hour timeframe, we speculate that the AHR is transiently activated. Of note, this temporary increase in KP metabolites stands in contrast to the permanent exposition to elevated AHR ligand availability that is frequently observed in various pathological conditions.

**Figure 2.**
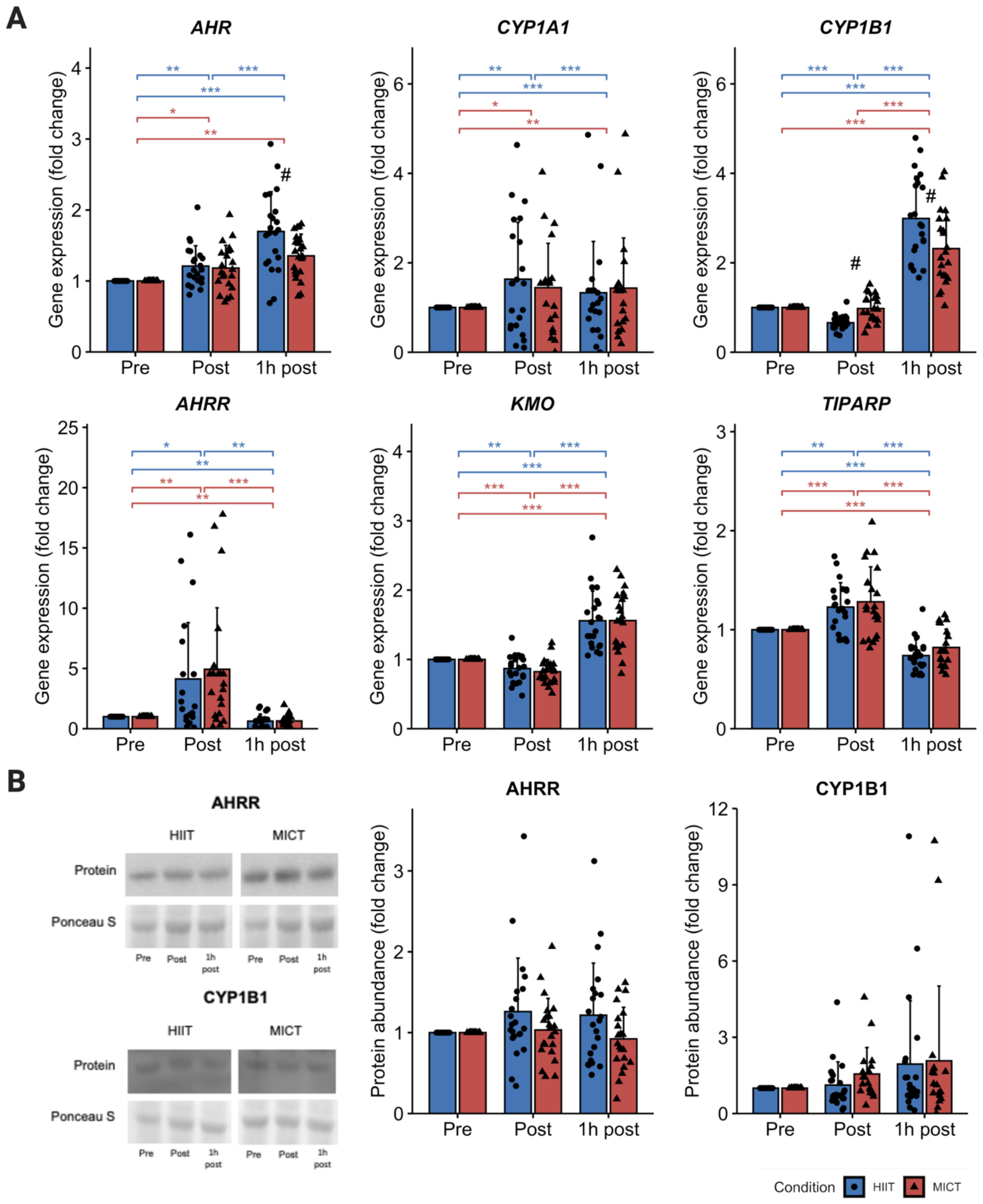
Acute exercise-induced AHR activation in PBMCs. A: Gene expression levels of AHR target genes in PBMCs in response to acute high-intensity interval training and moderate-intensity continuous training. Participants that could be evaluated: (for all outcome measures except *CYP1A1*) HIIT: n=22; MICT: n=22; *CYP1A1*: HIIT: n=21; MICT: n=22. B: Protein levels of key AHR targets in PBMCs in response to acute exercise. Participants that could be evaluated: AHRR: HIIT: n= 22; MICT: n=21, CYP1B1: HIIT: n=22; MICT: n=17. Significances are based on analyses of variance (ANOVAs) with repeated measures and Bonferroni-corrected post hoc comparisons. * p < .05; ** p < .001; *** p < .0001.

In conclusion, we identified the KP-AHR axis to be responsive to acute exercise in circulating immune cells. Considering the broad relevance of AHR signaling in regulating immunity and the implications for different diseases, our findings uncover a novel mechanistic link that potentially mediates exercise-induced health benefits. The results emphasize the superiority of training with higher intensities to initiate immunomodulatory processes – even when exercise modes are matched for duration and energy demand. Future trials are warranted to examine exercise-induced AHR activation in distinct immune cell subsets, other tissues, as well as potential functional consequences.

## Methods

The study was approved by the local ethics committee of the German Sport University Cologne and prospectively registered in the German Clinical Trials Register (DRKS00017686).

### Study design and participants

In total, 24 healthy adults (n=12 female; n=12 male) participated in this randomized cross-over trial that was designed to compare the acute effects of HIIT versus MICT on the kynurenine pathway-AHR axis. Participants reported to be recreational runner to ensure the achievement of exhaustion on the treadmill based cardiopulmonary exercise test and thus accurate controlment of training intensity during the HIIT and MICT bout, respectively (see Baseline assessment). All participants had a body mass index < 30 (kg×m^-2^) and were free of drug and supplement intake. Detailed participant characteristics are provided in table 1. Participants completed three laboratory visits at the German Sport University Cologne in this study. Participants refrained from any nutritional intake during the morning hours before each visit. Laboratory visits were all conducted at similar time of day between 07:00-10:00 AM in order to control for potential circadian influences. Participants refrained from caffeine and alcohol consumption 24h before each visit, and were allowed to drink water ad libitum during all three laboratory visits.

### Cardiopulmonary exercise test

A cardiopulmonary exercise test was conducted on a motorized treadmill (Woodway ELG 90, Weil Ma Rhein, Germany) to determine participants’ 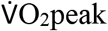 at an exercise physiology laboratory at the German Sport University Cologne. The cardiopulmonary exercise test was performed within four weeks prior to the interventions. Following a 5-minute warm-up period, increments of 1km^-1^ were applied until exhaustion. The starting speed for each test was set at 8km^-1^ and 1% incline was used for all procedures. Cortex Metalyzer 3B (CORTEX Biophysik GmbH, Leipzig, Germany) was used as spirometer and 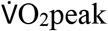 was determined by breath-by-breath analysis.

### Interventions and blood sampling

At least two days of rest were implemented after the cardiopulmonary exercise test. Participants then visited the exercise physiology laboratory in the morning between 07:00-10:00 AM for performing a single session of HIIT and MICT on separate days and similar time of day, respectively, with a wash-out period of at least seven days in between both sessions. The sequence of the HIIT and MICT session was randomized with concealed allocation by the software Randomization in Treatment Arms (RITA; Evident, Lübeck, Germany). Using an established and previously published running protocol^20,21^, HIIT and MICT sessions were matched for both energy demand and duration in order to isolate exercise intensity as sole diverging factor between both interventions. The HIIT session consisted of six 3-minute intervals at a speed corresponding to 90%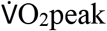. Between the intervals, 3-minutes of active recovery at a speed corresponding to 50% 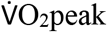 were applied. The HIIT session was framed by a 7-minute warm-up and cool down period at a speed corresponding to 70% 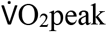, respectively. The MICT session was of equal duration (50 min.) and participants continuously exercised at a speed corresponding to 70% 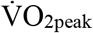.

Blood samples were collected from a median cubital vein at baseline (resting condition before exercise), immediately post exercise, and 1h post exercise at the HIIT and MICT session, respectively.

### Analytic procedures

#### Serum and PBMC isolation

Blood samples were collected in EDTA and serum tubes. Serum samples were centrifugated after 15min of clotting at 2500g for 10 minutes. Serum was aliquoted and stored at -80°C until HPLC MS/MS analysis. EDTA samples were diluted with PBS and carefully layered on top of a lymphocyte separation medium. After centrifugation, the PBMC containing layer was separated and washed with PBS. Cells were frozen at -150°C until further analyses.

#### Preparation of cell lysates

PBMCs were thawed, pelleted and resuspended in a lysis buffer composed of RIPA buffer, 10nM NaF, 1mM Na_3_VO_4_, Complete EDTA free protease inhibitor (Merck), Phospho Stop (Merck), 250U/ml Benzonase and 10U/ml DNase 1. Cells were lysed for 1h on ice and centrifuged at 13.000xg for 15 Min. Thereafter, the supernatant was transferred to a new tube and protein concentration was measured using the Pierce™ BCA Protein Assay Kit (Thermo Fischer). Cell lysates were stored at -80°C until analysis (western blotting).

#### HPLC MS/MS

Samples were thawed and 50µl of the serum were mixed with an internal standard and ice-cold methanol for protein precipitation. After centrifugation at 17.000xg at 4°C for 5 min, 100 µL of the supernatant was transferred into a glass vial for liquid chromatographic-mass spectrometric analysis (LC-MS/MS). For this purpose, a Waters ACQUITY UPLC® system interfaced via electrospray ionization (ESI) to a Xevo® TQ-XS triple quadrupole mass spectrometer (Waters, Eschborn, Germany) was employed. For separation an ACQUITY UPLC® HSS T3 analytical column (2.1 x 50 mm, 1.8 µm particle size) was used, with 5 mM ammonium acetate solution adjusted to pH 9 by adding NH_4_OH as aqueous solvent A and acetonitrile acidified with 0.1% formic acid as organic solvent B for Elution. The Elution was performed with a gradient of 2%-20% solvent B within 6 min followed by a steep increase to 100% within 1 min and held for 2 min. The ESI source was operated in positive mode using an impactor voltage of 3.5 kV and a desolvation temperature of 350 °C. For quantification and identification purposes, 2 diagnostic ion transitions (precursor ion to product ion) were used, which were recorded in multiple reaction monitoring (MRM) experiments. Argon (purity grade 5.0) was used for the collision-induced dissociations and collision energies were optimized for each analyte. A more detailed description was previously published elsewhere^22^.

#### Quantitative Real-Time-PCR (qRT-PCR)

RNA was isolated from PBMCs using a column-based isolation kit (Promega GmbH, Germany). RNA concentration and purity was measured on a Nanodrop™ microvolume spectralphotometer (Thermo Fisher Scientific, USA) and directly used for cDNA synthesis (Promega GmbH, Germany). qRT-PCR was performed for *AhR, AhRR, CYP1A1, CYP1B1, IDO1, IDO2, IL4I1, KAT3, KAT4, KMO*, and *TIPARP* in relation to the reference genes *ACTB, RPS18* and *UBE2D2* using the corresponding PCR master mix (Promega GmbH, Germany) on a CFX96 Detection System (Bio-Rad Laboratories GmbH, Germany). Primers were either published before or designed to span exon-exon-junctions. Primer details are shown in table S3 (supplementary material). Gene expression was calculated using the ΔΔCt method under consideration of PCR efficiency. Missing values and outliers defined as z score ≥ 3 (2.8% and 3.3%, respectively) were imputed with the mean value. All qRT-PCR data are reported as fold change relative to baseline.

#### Western Blotting

Before analysis, the PBMC lysates were diluted to a protein concentration of 2 mg/ml, suspended in 4x Laemmli buffer (#1610747; BioRad Laboratories GmbH, Munich, Germany) and heated at 95°C for 1 min.

Equal amounts of protein (15 μg) for each subject and time point were loaded on a 26-well (4– 12% BIS-TRIS Gel). Electrophoretic separation was conducted using a gel-casting system in MOPS electrophoresis buffer (all BioRad Laboratories GmbH, Munich, Germany). After electrophoresis, the gel was transferred to a polyvinylidene difluoride (PVDF) membrane (GE Healthcare Life Science, Amersham, UK) by semidry blotting (Trans Blot Turbo, Bio-Rad, Hercules, CA, USA) for 40 min (1.2 A, 25 Vmax). Equal sample loading and transfer were checked by staining the PVDF membrane with Ponceau S after transfer.

Membranes were then blocked for one hour at room temperature (RT) in 5% nonfat dry milk dissolved in tris-buffered saline supplemented with 0.1% Tween20 (TBST). Hereafter, primary antibodies were dissolved in 5% bovine serum albumin in TBST and incubated overnight for 17 hours at 4 °C. Afterwards, membranes were washed with TBST and then incubated with secondary antibodies diluted in TBST containing 5% nonfat dry milk for one hour at RT. After a series of washing steps, membranes were finally incubated for 3 min with an enhanced chemiluminescence assay (ECL-Kit, GE Healthcare Life Science, Amersham, UK) and digital images of the responsive bands automatically captured via an imaging system (ChemiDoc MP, Bio-Rad, Hercules, CA, USA). Subsequently, the membranes were stripped for 6 minutes at RT using a Restore Western Blot Stripping Buffer (#21059; Thermo Scientific, Waltham, MA, USA), blocked, incubated with primary antibodies for the detection of proteins with a different molecular weight and processed as described above.

Band densities were assessed semi-quantitatively using the ImageJ software (v. 1.53t; National Institute of Health, New York, NY, USA). The ratio of densities between each band and Ponceau S was calculated at all timepoints. All data is reported as fold change relative to baseline.

Antibodies used for western blotting: Primary: AHRR (rabbit polyclonal IgG AB; #PA5-107054 ; 1:250; Invitrogen, Rockford, IL, USA); CYP1B1 (rabbit polyclonal igG AB; #PA5-95277; 1:1000; Invitrogen, Rockford, IL, USA); secondary: anti-rabbit IgG, HRP-linked (#7074S; 1:7500; Cell Signaling Technology, Danvers, MA, USA); anti-mouse IgG, HRP-linked (#7076S; 1:7500; Cell Signaling Technology, Danvers, MA, USA).

### Statistical analyses

Statistical within and between group effects were evaluated using univariate analysis of variance models (ANOVA). Greenhouse-Geisser correction was used if sphericity was violated (Mauchly’s test). In case of statistical main effects of the ANOVA model, Bonferroni-corrected post hoc comparisons were applied (and illustrated in the figures). Area under the curve (AUC) was compared for systemic levels of tryptophan and its metabolites KYN, KA, and quinolinic acid using independent t-tests. Level of significance was set at alpha = 5%. Statistical analyses were conducted using IBM SPSS statistics version 29.

## Data availability

Raw data will be provided by the corresponding author upon reasonable request.

## Acknowledgements

We would like to thank all participants for their participation in this study.

