## Supplementary Material for "Acute exercise increases systemic kynurenine pathway metabolites and activates the AHR in human PBMCs"

**Supplement 1 (S1).** Ratios of kynurenine pathway metabolite levels in response to acute high-intensity interval training (HIIT) and moderate continuous intensity training (MICT) in healthy adults (n=24). Blue line = HIIT; Red line = MICT.

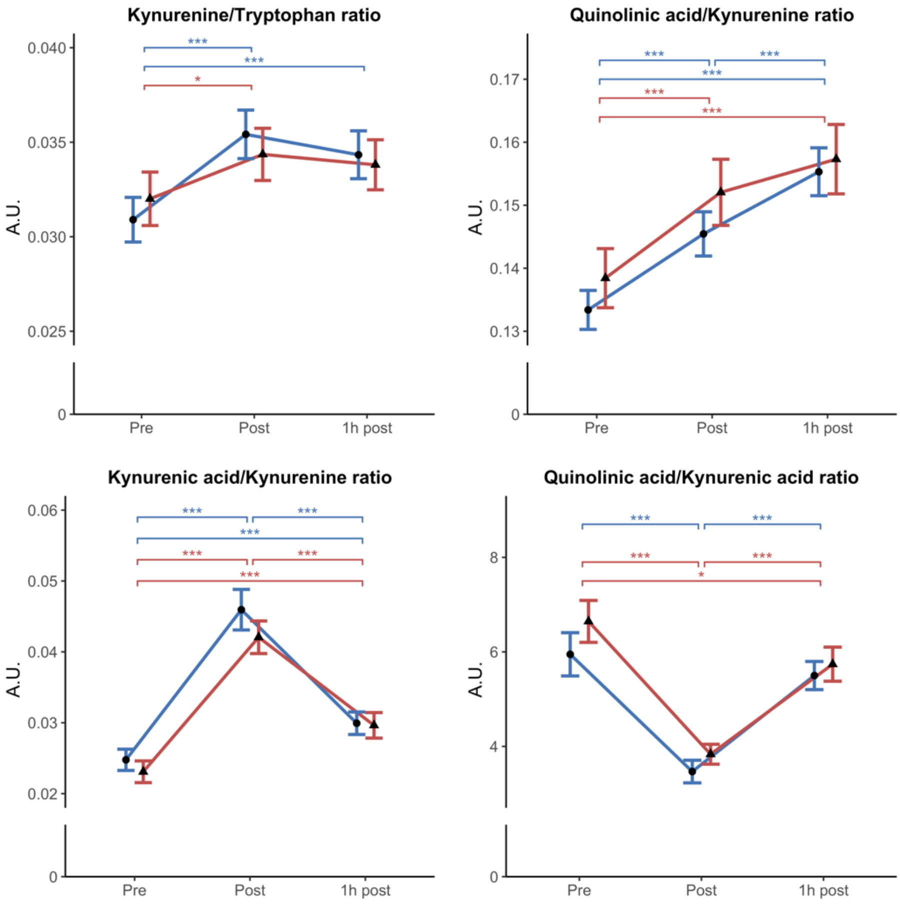

**Supplement 2 (S2).** Individual kinetics of gene expression levels of kynurenine pathway enzymes and AHR target genes in PBMCs in response to acute exercise. HIIT = high-intensity interval training. MICT = moderate-intensity continuous training. T0 = baseline; T1 = immediately post exercise; T2 = 1h post exercise.

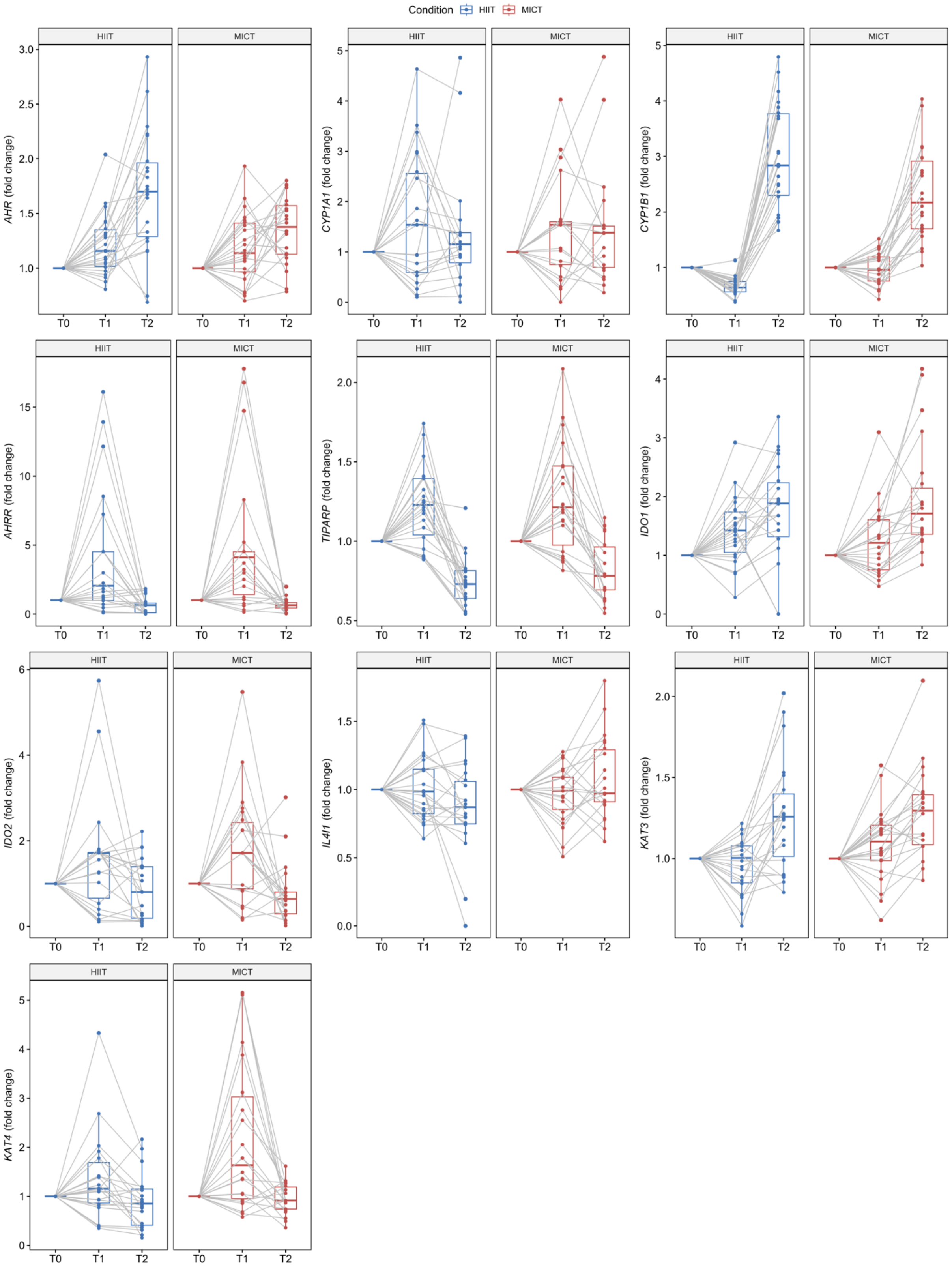

**Supplement 3 (S3).** Detailed ANOVA results comparing within- and between-group (HIIT vs. MICT) effects for all outcome measures. HIIT = high-intensity interval training. MICT = moderate-intensity continuous training. T0 = baseline; T1 = immediately post exercise; T2 = 1h post exercise.

| Measure | *df*$_1$ | *df*$_2$ | Time ($T_0$, $T_1$, $T_2$) | | Time*Group (HIIT vs. MICT) | |
|---|---|---|---|---|---|---|
| | | | *F* | *p* | *F* | *p* |
| Tryptophan | 2 | 90 | 2.08 | .131 | 0.25 | .248 |
| Kynurenine | 2 | 90 | 9.75 | <.001*** | 0.31 | .737 |
| Kynurenic acid | 1.31 | 56.30 | 105.83 | <.001*** | 1.67 | .203 |
| Quinolinic acid | 2 | 90 | 45.99 | <.001*** | 1.30 | .277 |
| *IDO1* | 1.67 | 70.13 | 27.22 | <.001*** | 1.15 | .316 |
| *IDO2* | 1.33 | 55.87 | 13.92 | <.001*** | 0.50 | .537 |
| *IL4I1* | 1.75 | 73.40 | 0.22 | .771 | 5.43 | .009** |
| *KAT3* | 1.66 | 69.83 | 29.79 | <.001*** | 1.60 | .212 |
| *KAT4* | 1.19 | 50.01 | 18.45 | <.001*** | 4.47 | .033* |
| *AHR* | 1.55 | 65.05 | 33.40 | <.001*** | 4.33 | .025* |
| *CYP1A1* | 2 | 82 | 4.54 | .014* | 0.25 | .712 |
| *CYP1B1* | 1.11 | 46.46 | 163.00 | <.001*** | 10.18 | .002** |
| *AHRR* | 1.02 | 42.73 | 25.46 | <.001*** | 0.29 | .596 |
| *KMO* | 1.31 | 54.94 | 99.76 | <.001*** | 0.13 | .787 |
| *TIPARP* | 1.51 | 63.49 | 74.64 | <.001*** | 0.565 | .524 |
| AHRR | 2 | 82 | 1.44 | .243 | 1.66 | .196 |
| CYP1B1 | 1.11 | 40.95 | 5.28 | .024* | 0.21 | .672 |

**p* < .05. ***p* < .01. ****p* < .001.

**Supplement 4 (S4).** Primers used for qRT-PCR.

| Target | Forward Primer | Reverse Primer | References |
|---|---|---|---|
| *ACTB* | GAGCTACGAGCTGCCTGACG | GTAGTTTCGTGGATGCCACAGG ACT | PMID: 16257224 |
| *RPS18* | CACTGCCATTAAGGGTGTGGG | CGTTCCACCTCATCCTCAGTG | |
| *UBE2D 2* | GTACTCTTGTCCATCTGTTCTCTG | CCATTCCCGAGCTATTCTGTT | PMID: 320051 48 |
| *AhR* | TAACCCAGACCAGATTCCTCCAGA | CCCTTGGAAATTCATTGCCAGA | PMID: 29320557 |
| *AhRR* | TGGCCTCTGGGCATTTATGG | GCTGGGCACTCGGTTAGAAT | |
| *CYP1A 1* | ACCCAGCTGACTTCATCCCTA | TGCTCAATCAGGCTGTCTGT | |
| *CYP1B 1* | GACGCCTTTATCCTCTCTGCG | ACGACCTGATCCAATTCTGCC | PMID: 219760 23 |
| *IDO1* | AATCCACAGGAAAATCTACCTGA T | AGCATGTTTAACTTCTCAACTCT TTCT | |
| *IDO2* | TGCGGAGCTATCACATCACC | GATGGTTTGGCTTCCCATGC | |
| *IL4I1* | TCTCAACCAGGCCCTCAAAG | GAGAAGATATTCCAAGAGCGTG T | |
| *KAT3* | CTGCAGACCCTTCTGTTGTGA | CAAGTGATGGATGGCCAAAGC | |
| *KAT4* | TTATATGGTGAGCGTGTAGGAGC | ACTTTCACTTCTTGCAGCCATT | |
| *KMO* | GGATCTATTGACTGCTGCTGAGAA ATAC | CCATCACATCCTACAATGAGGT CAC | |
| *TIPAR P* | CACCCTCTAGCAATGTCAACTC | CAGACTCGGGATACTCTCTCC | PMID: 219760 23 |
